# Semaglutide Alters Behaviour and Nucleus Accumbens Oscillatory Activity in Healthy Mice

**DOI:** 10.1101/2024.09.06.611514

**Authors:** Alejo Mosqueira, Sanaz Ansarifar, Sadegh Nabavi, Andrea Moreno

## Abstract

Semaglutide, a GLP-1 receptor agonist, is widely studied in metabolic disorders, yet its effects in healthy subjects remain unclear. Here, we show that daily administration of semaglutide (0.1 mg/kg) in healthy mice alters behaviour in various tests related to stress and reward pursuit. Additionally, acute administration of semaglutide changes the oscillatory activity in the nucleus accumbens in the delta, theta and alpha bands. These findings reveal that semaglutide impacts both behaviour and neural dynamics in non-diseased brains, offering a baseline for interpreting its broader therapeutic potential.

---

Semaglutide^1,2^, a long-acting glucagon-like peptide-1 (GLP-1) analogue traditionally used for the treatment of diabetes, recently emerged as an efficient therapeutic agent for obesity^3–6^. More recently, semaglutide has also been suggested as a promising treatment for addiction^7–10^, further expanding its potential applications beyond metabolic disorders. However, preclinical research has largely focused on obese and/or diabetic models, and its effects in healthy subjects remain largely underexplored^11–16^. This study aims to investigate the behavioural impacts of semaglutide in healthy C57BL/6JRj mice, providing insight into its broader neurobiological effects and establishing a baseline for future research in both metabolic and neuropsychiatric contexts.

Different administration protocols of semaglutide were tested on adult healthy male C57BL/6JRj wildtype mice, a commonly used strain in behavioural research. The first protocol consisted of subcutaneous injections of semaglutide (Ozempic®, 0.1 mg/kg) on a weekly basis for eight weeks. This regimen mirrors human clinical practice for chronic conditions^3^. To account for the higher metabolic rate in mice and aiming to explore short-term effects^17^, the second protocol consisted of daily injections of the same dose for eight consecutive days prior to the start of behavioural testing, followed by additional daily injections at the end of the test sessions. All mice underwent a series of behavioural tests designed to evaluate different aspects of behaviour and emotional state, which included open field test, marble burying test, nestlet shredding test, female urine sniffing test, and forced swimming test (Figure 1A). Throughout the study period, we monitored food and water consumption and body weight to assess any physiological changes. As previously reported in rodents ^8,9,18–20^, we observed a substantial weight loss after only a single injection that continued throughout the treatment period (Figure 1B). Notably, food and water consumption were also reduced (Figure 1B), which may have contributed to the observed weight loss. Below, we report the behavioural changes associated with the daily injection treatment.

**Figure 1.**
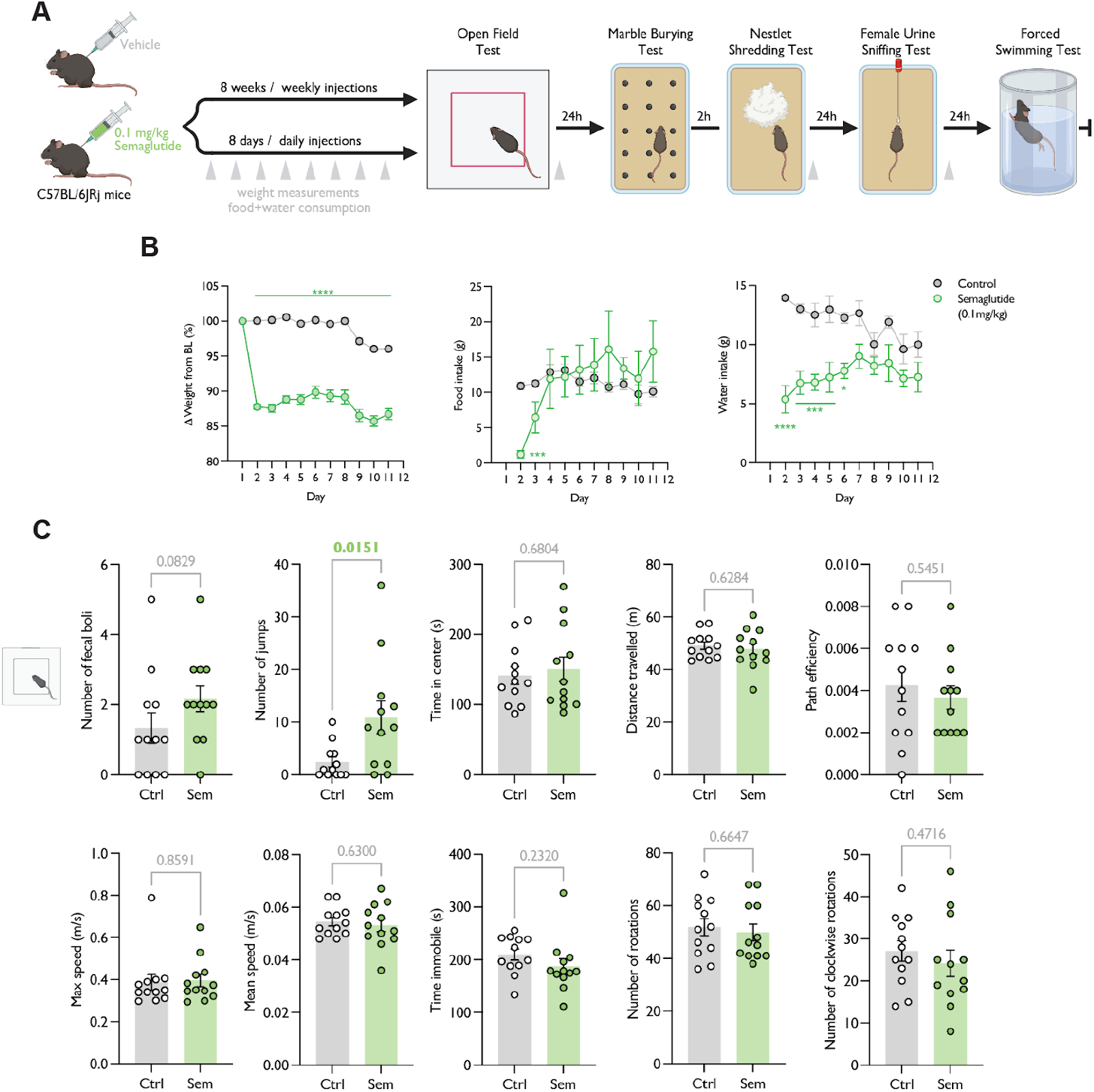
Daily semaglutide administration in healthy mice causes significant weight loss and increases jumping behaviour in the open field test. (A) Experimental design (see Methods). (B) Daily weight difference, food and water consumption. Weight is measured per animal (n=12 per group); food and water consumption are measured per cage as animals are group-housed, with 3 animals per cage (n=4 per group). (C) Open Field Test results. No statistical differences were detected in any of the tested variables, with the exception of jumping behaviour, which was higher in the treated group. N=12 per group; Mean±SEM; p-values per comparison (statistically significant p-values are highlighted in bold and green).

The open field test, a standard measure of locomotion and anxiety, revealed no changes in any of the measured variables (faecal boli, time spent in the field’s centre, distance travelled, path efficiency, maximum speed, mean speed, time spent immobile, number of rotations, or number of clockwise rotations; Figure 1C). However, we observed a significant increase in jumping behaviour (Figure 1C), with the semaglutide group jumping significantly more than the control group (Median = 9 vs. 1; U = 31, p = 0.0151; Mann-Whitney U test). This could indicate more attempts to escape the testing box, as the jumping was most frequently observed towards the walls.

In order to evaluate anxiety and compulsive behaviour, we applied two complementary tests: the marble burying test and the nestlet shredding test ^21,22^. No significant changes were observed in the marble burying test (Figure 2A) between the groups (U = 44.50, p = 0.1157). In contrast, the nestlet shredding test (Figure 2B) demonstrated a significant reduction in shredding among the semaglutide-treated mice (Mean = 1.615) compared to controls (Mean = 6.187), with a difference of 4.572 ± 1.117 (t(22) = 4.095, p = 0.0005, R^2^ = 0.4325). This finding suggests an increase in anxiety or a shift in exploratory behaviour, potentially related to semaglutide’s impact on the central nervous system.

**Figure 2.**
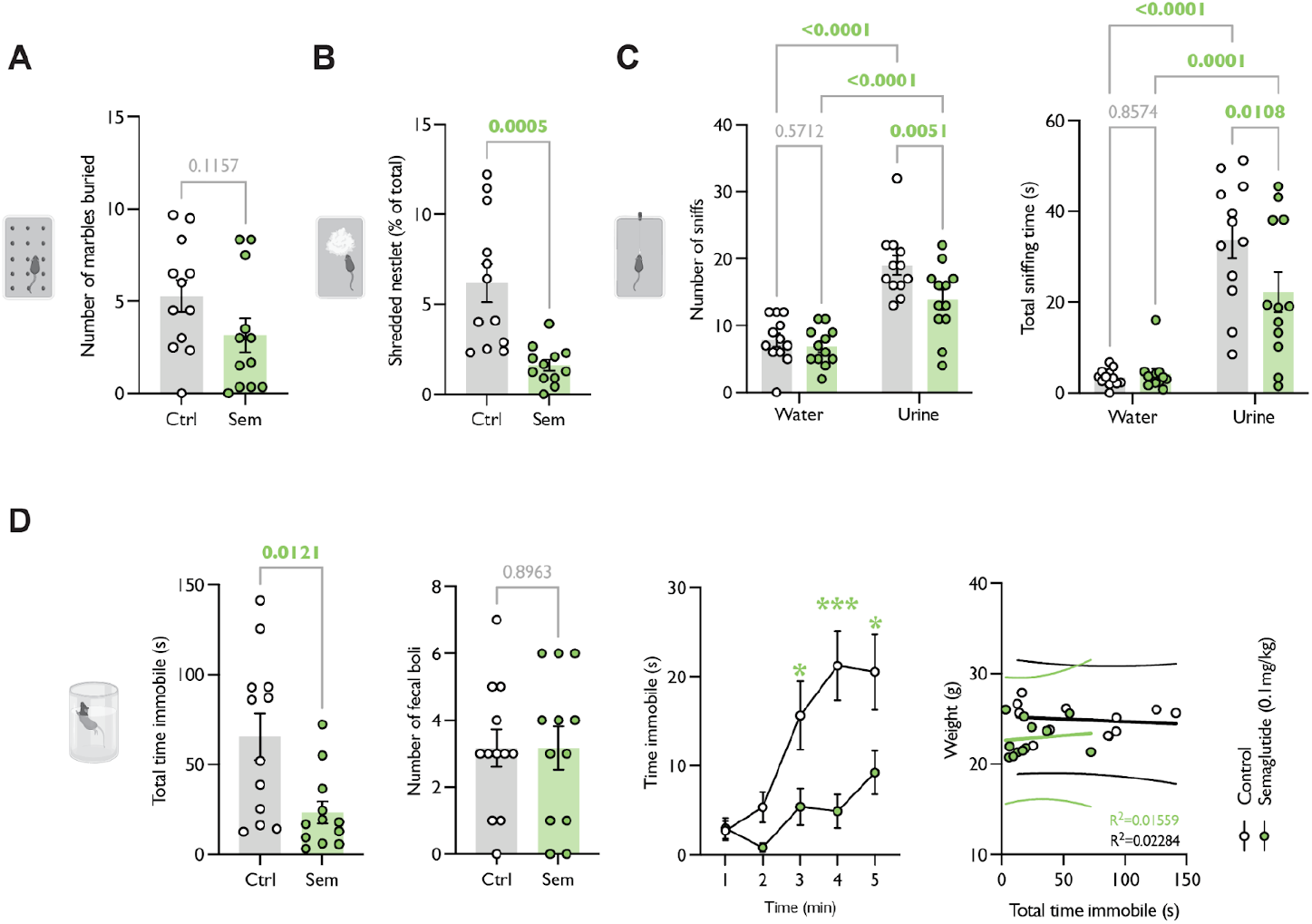
Daily semaglutide administration in healthy mice causes changes in the Female Urine Sniffing Test, Nestlet Shredding Test and Forced Swimming Test, but not in the Marble Burying Test. (A) Marble Burying Test results. (B) Nestlet Shredding Test results. (C) Female Urine Sniffing Test (FUST) results. (D) Forced Swimming Test results. In all graphs, n=12 per group; Mean±SEM; p-values per comparison (statistically significant p-values are highlighted in bold and green).

The Female Urine Sniffing Test (FUST) (Figure 2C) is used to evaluate reward-seeking behaviour in rodents^23^ by measuring the time spent sniffing cotton swabs soaked in different substances. We employed this test to assess reward-seeking as an independent metric, separate from food consumption variables that might be influenced by the primary effects of semaglutide. First, the rodent is exposed to a swab soaked in water, followed by one soaked in urine from females in the estrous cycle. Healthy male rodents typically spend significantly more time sniffing the female urine, indicating natural reward-seeking behaviour. A decrease in this behaviour is linked to anhedonia. Both groups showed a significant overall difference between sniffing water and urine (F(1, 22) = 81.94, p < 0.0001 for number of sniffs; F(1, 22) = 76.16, p < 0.0001 for sniffing time). A two-way repeated measures ANOVA revealed a significant interaction between scent sample and treatment (F(1, 22) = 4.310, p = 0.0498), indicating that semaglutide treatment affected sniffing behaviour differently, depending on the olfactory cue presented. Specifically, the number of sniffs directed at urine was significantly reduced in the semaglutide-treated group compared to controls (Semaglutide: 13.83 sniffs, Control: 19 sniffs; p = 0.0051, total reduction of 27.21%). Total sniffing time also showed a significant smell x treatment interaction (F(1, 22) = 4.881, p = 0.0379), with the semaglutide-treated group spending significantly less time sniffing urine, representing a 33.9% reduction (Semaglutide: 22.18 seconds, Control: 33.6 seconds; p = 0.0108).

In the forced swimming test (Figure 2D), a common method for assessing depressive-like behaviour and resilience^24^, the semaglutide-treated group exhibited a 74.8% reduction in immobility time. (Median = 17.40 vs. 69.05, p = 0.0121, Mann-Whitney U = 29). Additionally, when examining immobility per minute, a two-way repeated measures ANOVA showed a significant effect of treatment over time (F(1, 22) = 8.643, p = 0.0076), with semaglutide-treated mice spending significantly less time immobile in minutes 3, 4, and 5 compared to controls (p values of 0.0309, 0.0001, and 0.0131, respectively). Within-group linear regression analyses revealed no significant association between body weight and total time immobile in either the control (R^2^ = 0.02284, p = 0.6392) or semaglutide-treated group (R^2^ = 0.01559, p = 0.6990), indicating that the observed immobility differences are unlikely to be confounded by body weight. The overall reduction in immobility points to a potential improvement in coping behaviour or a reduction in depressive-like responses.

In contrast to the daily-injected group, the weekly-injection group did not exhibit significant behavioural changes. This lack of effect highlights the potential influence of dosing frequency on the behavioural outcomes associated with semaglutide. It shows that the daily dosing regimen is more likely to elicit behavioural changes, which could be due to differences in drug metabolism or pharmacodynamics, or short-term side effects. Despite the absence of significant behavioural changes, the weekly injection regimen was still effective in promoting weight loss. This indicates that while dosing frequency may influence behaviour, the metabolic effects of semaglutide on weight reduction remain robust even with less frequent administration (see Supplementary Figure 1).

We next performed local field potential (LFP) recordings from the nucleus accumbens under anaesthesia before and after acute treatment (Figure 3). By acute treatment, we refer to a single administration of semaglutide and the immediate effects within the same recording session, as opposed to repeated or chronic dosing paradigms. We targeted the nucleus accumbens to assess whether semaglutide modulates neural activity in a region that plays a well-established role in reward processing (potentially underlying the behavioural changes observed in this study), yet relatively understudied in this context. We examined both aperiodic and periodic components of the LFP^25^. Aperiodic components represent non-rhythmic, baseline neural activity, indicating general neural network states. Periodic components, like theta oscillations, reflect synchronised neural activity linked to specific functions such as motivation and reward.

**Figure 3.**
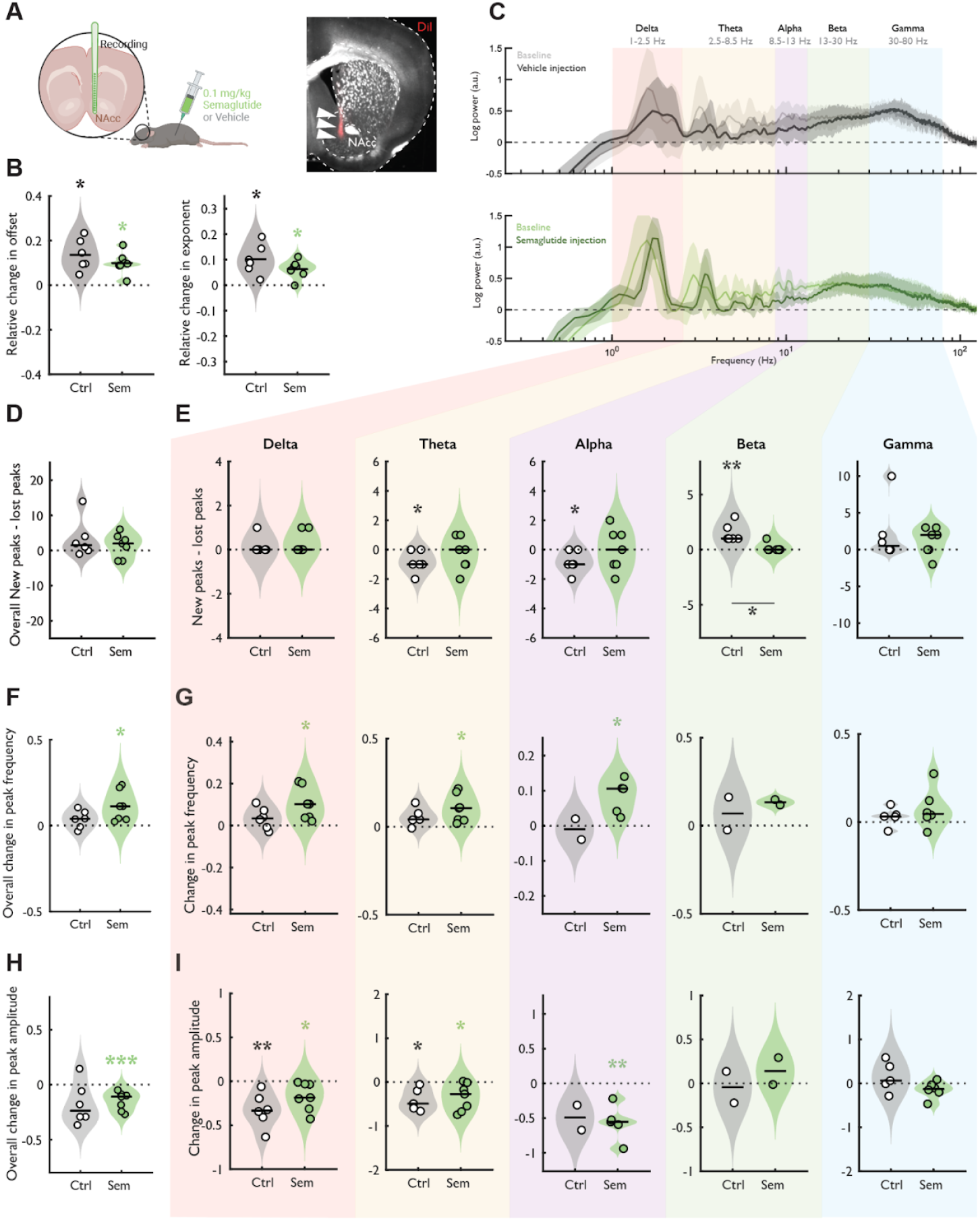
A single semaglutide administration in anaesthetised healthy mice alters oscillatory activity recorded in the nucleus accumbens. (A) Experimental design. (B) Changes in aperiodic components (offset and exponent), relative to baseline, for each treatment. (C) Baseline vs. post-injection spectrum (after subtraction of aperiodic component) for vehicle (top) and semaglutide (bottom). (D) Overall number of new vs. lost peaks. (E) New vs. lost peaks by frequency band. (F) Overall frequency shift. (G) Frequency shift by frequency band. (H) Overall changes in peak amplitude. (I) Changes in peak amplitude by frequency band. In all graphs, n=6 for the Control group and n=7 for the Semaglutide group; statistical significance is shown as comparisons against baseline; * = p≤0.01; ** = p≤0.05; *** = p≤0.005.

Analysis of the aperiodic component (Figure 3B) revealed a significant increase in the offset and exponent for both the control (offset: p=0.005, slope: p=0.008, one-sample t-test) and semaglutide group (offset: p=0.002, slope: p=0.003, one-sample t-test). However, there were no statistical differences between the groups (offset change: 0.14±0.07 vs 0.10±0.05, Ctrl vs Sem, p=0.28, two-sample t-test; exponent change: 0.10±0.06 vs 0.06±0.04, Ctrl vs Sem, p=0.17, two-sample t-test). This suggests that semaglutide did not induce global changes in background neural activity.

To study the impact of semaglutide on narrowband LFP oscillations, we analysed changes in the frequency components or periodic components of the signal (Figure 3C). First, we calculated the difference in the number of newly detected peaks and lost peaks after injection compared to baseline measurements (Figure 3D-E; ‘New-lost peaks’, see Methods). In this case, positive values indicate that more peaks were detected following the injection than at baseline. We did not see any differences in this parameter globally (Figure 3D). However, we did see a significant decrease in detected peaks in the theta (p=0.04, one-sample t-test) and alpha (p=0.04, one-sample t-test) bands, and a significant increase in detected peaks in the beta (p=0.007, one-sample t-test) band, but only in the control group (Figure 3E). This indicates that vehicle injection alone modulated peak detectability in theta, alpha, and beta bands, which may indicate that the injection process (e.g., minor fluctuations in brain state due to handling, fluid pressure changes, etc.) slightly altered the brain’s oscillatory activity or the ability of our method to detect them. However, since we did not observe this in the semaglutide group, we did not consider this a reason for exclusion.

Second, we observed that semaglutide significantly promoted an overall peak frequency shift relative to baseline (Figure 3F; 0.11 ± 0.09, p = 0.015, one-sample t-test), indicating a shift in the dominant oscillatory components towards higher frequencies. To better understand which components were driving this effect, we analysed frequency shift by band (Figure 3G). This revealed that the increase was primarily driven by significant frequency shifts toward higher frequencies in the delta (p=0.013, one-sample t-test), theta (p=0.013, one-sample t-test) and alpha (p=0.019, one-sample t-test) bands, suggesting that semaglutide selectively affects slower oscillatory activity. In contrast, no significant frequency shifts were observed in the control group, either globally or within specific bands, highlighting a treatment-specific spectral modulation by semaglutide.

In parallel, semaglutide also significantly reduced overall peak amplitude compared to baseline (Figure 3H; −0.15 ± 0.08, p = 0.004, one-sample t-test), indicating a decrease in the strength or power of dominant oscillatory components. Band-specific analysis (Figure 3I) revealed that this reduction was significant in the delta (p=0.03, one-sample t-test), theta (p=0.03, one-sample t-test) and alpha (p=0.008, one-sample t-test) bands. These changes suggest a broad attenuation of low-frequency oscillatory power following semaglutide administration. In the control group, reductions in peak amplitude were observed only in the delta (p = 0.009) and theta (p = 0.03) bands, with no significant changes in alpha, again suggesting a more robust and widespread effect of semaglutide on spectral power.

These findings suggest that semaglutide alters delta, theta and alpha oscillatory patterns, specifically promoting a shift towards higher peak frequencies. Since theta oscillations play a key role in motivated behaviour, if the observations in anaesthetised subjects are translated to awake ones, an increased theta frequency could relate to heightened locomotion and reduced immobility (both of which were observed in the behavioural tests), reflecting enhanced arousal ^26–28^, or changes in the mechanisms governing action selection and behavioural inhibition^29^. Since anaesthesia affects network activity, future studies could examine semaglutide’s effects on theta oscillations in awake, freely moving animals and employ source localisation methods to determine whether these changes arise from local accumbal activity or broader circuit-level interactions.

Overall, our results are aligned with previous reports of semaglutide reducing the pursuit of rewards, although previous literature has mainly focused on addiction-related rewards (such as alcohol ^7–10,19,30^ and cocaine ^31^). Our electrophysiological findings support others that report semaglutide’s access to the nucleus accumbens in vivo after a systemic injection ^9^. These results could pave the way to a better understanding of semaglutide’s effects on the reward processing systems in healthy subjects, where addiction-related brain circuit changes are not present.

Another notable finding was the significant reduction in immobility time during the forced swimming test in mice treated with daily semaglutide injections, a behavioural change typically interpreted as indicative of antidepressant effect. Although previous studies investigating the relationship between semaglutide and depressive symptoms have reported mixed findings ^32–34^, these studies are usually focused on patients or animal models of obesity and/or diabetes. Our results provide evidence suggesting a potential protective effect of semaglutide against depression-like or stress-induced behaviours in healthy animals.

While we did not include a weight-matched control group, such as mice on a calorie-restricted diet, the results from the group that received weekly injections of semaglutide (who gained significantly less weight that the control group but showed no behavioural changes) and the absence of correlation between animal weight and time immobile in the FST test suggest that factors beyond weight differences may influence the observed outcomes. Notably, the most pronounced behavioural differences in this study, including reduced urine sniffing time and nestlet shredding, along with the electrophysiological changes, cannot be solely attributed to weight differences.

The dose of semaglutide used in this study (0.1 mg/kg) may have allowed for behavioural effects to be observed with daily dosing but not with weekly injections. This dose, which has been extensively used in rodent studies in order to match human plasma concentrations ^9,19,35,36^ was chosen to be conservative in assessing the drug’s impact on behaviour, but it is possible that a higher dose could reveal more consistent effects across different dosing regimens.

This study underscores the importance of considering both dosage and administration frequency in behavioural research, as different regimens can yield varying effects. The observed behavioural changes following daily semaglutide dosing could guide future research into its effects across various models and health conditions ^37,38^, such as obesity ^10,20^, neurodegenerative diseases ^39–41^, mood disorders, or addiction ^9,19^. In conclusion, daily administration of semaglutide in healthy male C57BL/6JRj mice results in notable behavioural effects, including increased jumping activity, reduced nestlet shredding, decreased female urine sniffing, and lower immobility in the forced swimming test. A single dose of semaglutide induces alterations in nucleus accumbens LFP activity, particularly in the delta and theta range, where semaglutide increased their frequency. As theta rhythms play a crucial role in motivated behaviour, these findings suggest that semaglutide’s effects extend beyond metabolic regulation to influence neural circuits that could be involved in motivation and reward processing.

## Methods

### Animals

Male C57BL/JRj mice were purchased from Janvier Labs at 7 weeks old, and used in experiments at 8-18 weeks of age. Mice were group housed in cages of 3, and provided with ad libitum access to food (regular chow) and water. Mice were housed under regular lighting conditions (12h light cycle, lights on at 6:00 AM). All injections, weighing procedures, and behavioural experiments were performed between 8:00 AM and 2:00 PM.

### Behaviour

Mice were split into weekly and daily groups. The weekly group received 8 injections (1 per week) of either vehicle or semaglutide, and weight/food/water measurements every week for 8 weeks, and were tested after week 8 on the described behavioural battery. The daily group received 8 injections (1 per day) for a week, while weight/food/water measurements were also performed daily. The daily group was then tested after day 8 in the behavioural battery and continued being given injections every day for 3 extra days while being tested. Overall, the daily group received 11 injections by the end of the behavioural battery timeline. Injections and weight/food/water measurements on the behaviour testing days were performed after behaviour tests to avoid short-term effects on behaviour due to the injections and to minimise animal manipulation. All mice (weekly and daily groups) started behavioural testing after 8 injections.

#### Open Field Test

Mice were placed in an arena (45 x 45 x 45 cm) made of polycarbonate with a paper flooring (30% Ethanol was used to clean between mice, and paper flooring renovated for each mouse). The arena was placed inside a noise-cancelling box (80 x 60 x 62 cm). Mice were allowed to explore the arena for 15 min. Mice were tracked and standard measurements of speed, zone visits, distance travelled and other measures of locomotion were automatically scored by Anymaze software. Faecal boli were manually counted at the end of the test. Jumping was manually counted offline based on the recorded videos.

#### Marble Burying Test

Standard cages with flat lids, containing one layer of 5 cm of bedding, were used to position a grid of 5×4 standard glass toy marbles (blue, 15 mm diameter, 5.2 g) evenly spaced throughout the cage. Mice were individually placed in the cage and left undisturbed (cage inside of a noise-cancelling box) for 30 mins. After the test, mice were removed and cages were photographed to evaluate (manually and semi-automatically) the number of marbles buried. See Supplementary Material (Supplementary Figure 2) for a discussion on marble burying quantification.

#### Nestlet Shredding Test

Mice were individually placed in cages with flat lids containing a layer of bedding and one new cotton nestlet, and left undisturbed in the cage in a noise-isolated box for 30 min. Nestlets were weighed before and after the test; weight differences are reported as shredding %.

#### Female Urine Sniffing Test

In order to determine the estrous stage of the females used to harvest the urine samples used in the test, vaginal cell samples (25 µL, in sterile saline solution) were taken from females, stained with cresyl violet (0.1%), and assessed under the microscope to identify estrous stage. Urine from females (from the same genetic background as the males) in estrous was harvested on the test day. Samples of 40 µL of urine were pipetted into cotton swabs and kept in closed test tubes under a fume hood until the testing phase. Urine samples were taken from the same batch for all tested mice in one group (two groups of n=12 mice).

The test followed several phases as described in^23^, administered during one day: (1) Habituation to a cotton swab without any liquid on it for 1 h; (2) Exposure to a cotton swab with water for 3 min; (3) Waiting time without a cotton swab in the cage for 45 min; (4) Exposure to a cotton swab with urine for 3 min. In stages 2 and 4, the sniffing was measured manually based on video recordings of the mice’s behaviour performed with Anymaze with a camera placed above the cage.

Data for the weekly injection group was excluded from this test, as the control animals in this group did not perform as expected according to the test’s assumptions (which expect control animals to interact preferentially with the female urine in comparison to water)^23^. Consequently, the data was deemed non-valid and not included in the final analysis.

#### Forced Swimming Test

Mice were placed in cylindrical containers (30 cm tall, 17 cm diameter) containing room-temperature water (up to a measure of 15.5 cm from the bottom of the container) for 5 minutes. Mice were tracked automatically with a side-camera with Anymaze to detect immobility episodes. Mice were sacrificed after finishing the test.

### Electrophysiology

#### Recordings

Neural activity was recorded from the nucleus accumbens under anaesthesia to assess the effects of semaglutide on local field potentials (LFPs). Mice were briefly anaesthetised with isoflurane (1.5-2%) and then injected with a mixture of fentanyl (0.05 mg/kg), medetomidine (0.5 mg/kg) and midazolam (5 mg/kg) – FMM (10 ml/kg, i.p.) to maintain the anaesthesia plane during the duration of the recording. Anaesthesia was maintained at a stable level throughout the experiment without additional doses. Body temperature was kept constant at 37 °C using a feedback-regulated heating pad. A 32-channel silicon probe (Neuronexus, Ann Arbor, US) was lowered into the nucleus accumbens using a stereotaxic frame (coordinates: AP +1.9 mm from bregma, ML +0.7 mm from bregma, DV −4.4 mm from surface). Raw data were filtered (0.1–3000 Hz), amplified (100x), digitised and stored (25 kHz sampling rate) for offline analysis, with a tethered recording system (Multichannel Systems, Reutlingen, Germany). Baseline recordings were performed for 30 minutes, after which semaglutide (0.1 mg/kg, i.p.) or vehicle (saline solution, i.p.) was administered. Post-injection recordings were performed for 1 hour immediately after injection to capture changes in oscillatory activity. Following recordings, mice were sacrificed and brains were extracted and fixed in 10% formalin solution for 48-72 h. Histological verification of electrode placement was performed by obtaining 100 µm slices from the region of interest with a vibratome. Electrode tracks were marked with DiI and confirmed via fluorescent imaging, as shown in Supplementary Figure 3.

#### Analysis

Local field potentials (LFPs) were analysed using custom scripts, built-in MATLAB functions, and the FOOOF toolbox^25^. LFPs were first decimated to 1 kHz using MATLAB’s ‘decimate’ function. The power spectral density (PSD) was then estimated using Welch’s method (‘pwelch’, window = 5000, fs = 1 kHz) in two time 10-minute periods: baseline (starting 15 minutes prior to injection) and post-injection (starting 50 minutes after injection). To mitigate power line noise, we linearly interpolated the power spectra in a [-0.25, 0.25] Hz window around 50 Hz and its harmonics (‘interp1’ function). We then used the FOOOF toolbox to extract both aperiodic (offset and exponent) and periodic (peak frequency, width, and amplitude) components from the PSDs (FOOOF parameters: f_range = [0.2, 150], peak_threshold = 1.7, min_peak_height = 2, peak_width_limits = [0.2, 20]). To quantify frequency shifts, we removed the aperiodic component from both baseline and post-injection spectra, transformed the remaining data to a logarithmic scale, smoothed it using a Gaussian kernel (‘smoothdata’, method = ‘gaussian’, window = 20), and applied a cross-correlation analysis (‘xcorr’). The resulting shift was then used to match baseline to post-injection peaks. To do this, we shifted the baseline peaks’ frequencies and looked for the closest match in the post-injection period (within a 10% frequency tolerance). When two post-injection peaks matched the same baseline peak, only the one consistent with the overall frequency shift (i.e., the rightward peak for forward shifts and the leftward peak for backward shifts) was kept. The matched peaks were used to quantify relative changes (relative to baseline) in peak amplitude and frequency following injections. To assess whether there was a significant change in peak count post-injection, we defined a metric called ‘New-lost peaks.’ This metric is calculated by subtracting the number of ‘lost’ peaks (detected at baseline but not post-injection) from the number of ‘new’ peaks (detected post-injection but absent from baseline). A positive ‘New-lost peaks’ value indicates more peaks detected post-injection than at baseline. For frequency band-specific analyses, we defined the following intervals: delta (1-2.5 Hz), theta (2.5-8.5 Hz), alpha (8.5-13 Hz), beta (13-30 Hz), and gamma (30-80 Hz).

### Statistical Analyses

Statistical analyses were performed using GraphPad Prism 10. In bar graphs, data are represented as Mean ± SEM. Group comparisons were made using either unpaired t-tests or Mann-Whitney U tests, depending on the data distribution. For repeated measures, two-way repeated measures ANOVA was applied, followed by post-hoc tests where appropriate. Significance was set at p < 0.05. For all tests, assumptions such as normality and sphericity were assessed, and non-parametric tests were used if assumptions were violated.

## Author contributions

AM✉ conceived the idea, planned the experiments, did behavioural and electrophysiological experiments, analysed the data, wrote the manuscript and prepared the figures. SA* did behavioural and electrophysiological experiments. AM* did pilot electrophysiological experiments, performed data analysis for behaviour and electrophysiology data, and developed the marble counting semi-automatic method. All authors contributed to the experimental design planning. All authors read, commented on and approved the manuscript.

## Acknowledgements

This study was supported by PROMEMO (Center of Excellence for Proteins in Memory funded by the Danish National Research Foundation) funding to SN (DNRF133). We would like to acknowledge the technical support offered by Anne-Katrine Vestergaard, and all our colleagues who commented on the initial experimental plan, ideas and hypothesis.

