## Supplementary material for "Semaglutide Alters Behaviour and Nucleus Accumbens Oscillatory Activity in Healthy Mice"

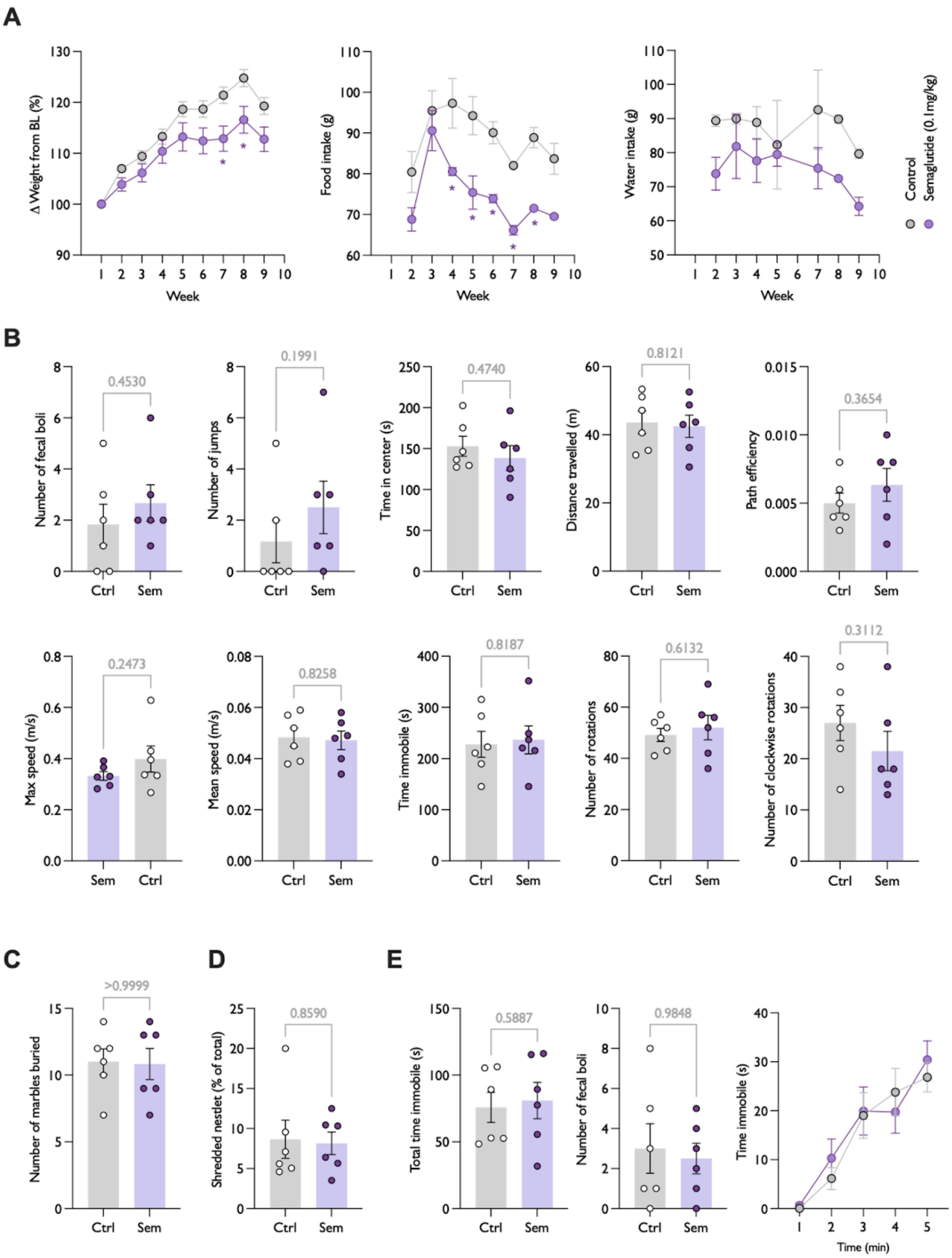


**Supplementary Figure 1. No behavioural effects were detected when administering Semaglutide on a weekly schedule.** (A) Weekly weight difference, food and water consumption. Weight is measured per animal (n=6 per group); food and water consumption are measured per cage as animals are group-housed, with 3 animals per cage (n=2 per group). (B) Open Field Test results. No statistical differences were detected in any of the tested variables. (C) Marble Burying Test results. No statistical differences were detected. (D) Nestlet Shredding Test results. No statistical differences were detected. (E) Forced Swimming Test results. No statistical differences were detected. In all graphs, n=6 per group; Mean±SEM; p-values per comparison (statistically significant p-values are highlighted in bold and purple).

**Supplementary Method 1. Semi-automatic marble burying quantification**

To quantify the marble-burying test, a custom semi-automated algorithm was developed in MATLAB (MathWorks, R2019b). Initially, the average size of a marble was estimated by manually creating circular regions of interest (ROIs) surrounding unburied marbles and then averaging the resulting ROIs' areas (Suppl. Figure 2A). This estimation was later used to quantify the number of marbles buried and normalise the total surface of unburied marbles, enabling comparison between different pictures. Subsequently, the area corresponding to the cage floor containing marbles was manually selected by creating a rectangular ROI (Suppl. Figure 2B). The algorithm was only applied to this ROI to prevent objects, reflections, or shadows outside the cage or on the cage's walls from being misclassified as marbles. A 2-D median filter was then applied to the ROI to mitigate the specular highlights on the marbles that could be identified as holes and sudden changes in contrast in the bedding material that could be falsely detected as marbles (Suppl. Figure 2C), followed by image binarisation through colour thresholding (Suppl. Figure 2D). The latter step was performed by inspecting the ROI for pixels where the three channels (R, G and B) were below a chosen threshold and replacing them with white values, whereas the remaining pixels were replaced with black values. The optimal neighbourhood size for the filtering step and the threshold for the image binarisation are dependent on the image size and the experimental setup (e.g., marble colour, bedding material, light conditions, etc.). Therefore, it is recommended to optimise these values by visually inspecting the results obtained for a few cases and fixing them for all images. In our case, optimal results were obtained with a neighbourhood size of 55 pixels and a threshold of 50 on a 0-255 scale. Morphological operations were then performed on the binary image to fill out holes resulting from specular highlights on the marbles and to correct for the underestimation of the marbles' edges resulting from the filtering step. This involved creating a disk-shaped structuring element with a radius of 10 pixels and applying morphological closing, followed by dilation to a new disk-shaped structuring element with a radius of 4 pixels. Finally, the marbles' boundaries were detected (Suppl. Figure 2E), and the number of marbles buried as well as the normalised marble buried area were computed (Suppl. Figure 2F). This was achieved through the calculation of the number of marbles with less than 1/3 of the area unburied (i.e., at least 2/3 of the area buried) and the subtraction of the area covered by unburied marbles (normalised to a single marble area) from the total number of marbles in the cage, respectively. The code will be made publicly available upon article publication.

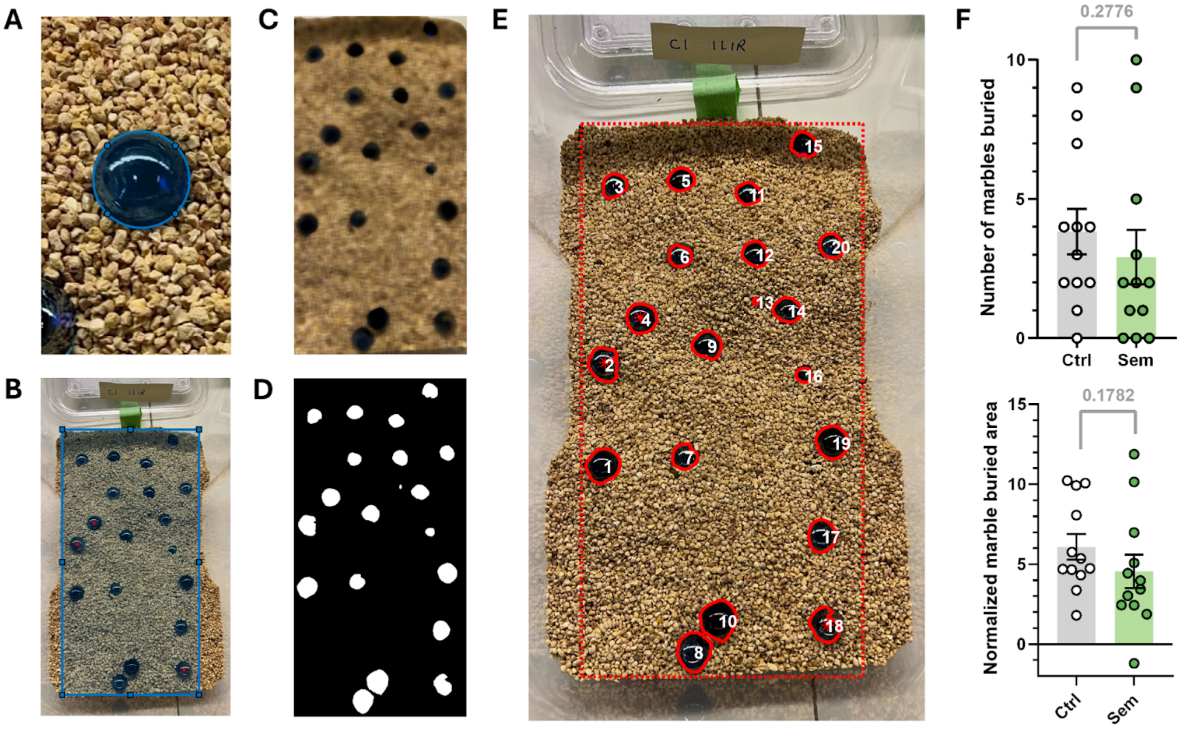


**Supplementary Figure 2. Semi-automated algorithm for quantification of the marble-burying test.** (A) Example of a circular ROI defined around a marble to estimate the average marble area. (B) Example of a rectangular ROI defined to delimit the region of the cage floor containing marbles. (C) A 2D median filter is applied to the ROI defined in B. This allows for the smoothing of the image of the bedding material and for the removal of specular highlights on the marbles. (D) Image binarisation of C. (D) Marbles' boundaries obtained from the binarised image shown in D (red solid lines around the marbles) are overlaid on the original picture. The analysed ROI is depicted by a red dashed rectangle. A unique number is assigned to each detected marble. Marbles used to estimate the average marble area are marked with an asterisk. (F) Results obtained for the semi-automatic quantification of the marble-burying test. The statistics show the p-values obtained by applying a Student's t-test.

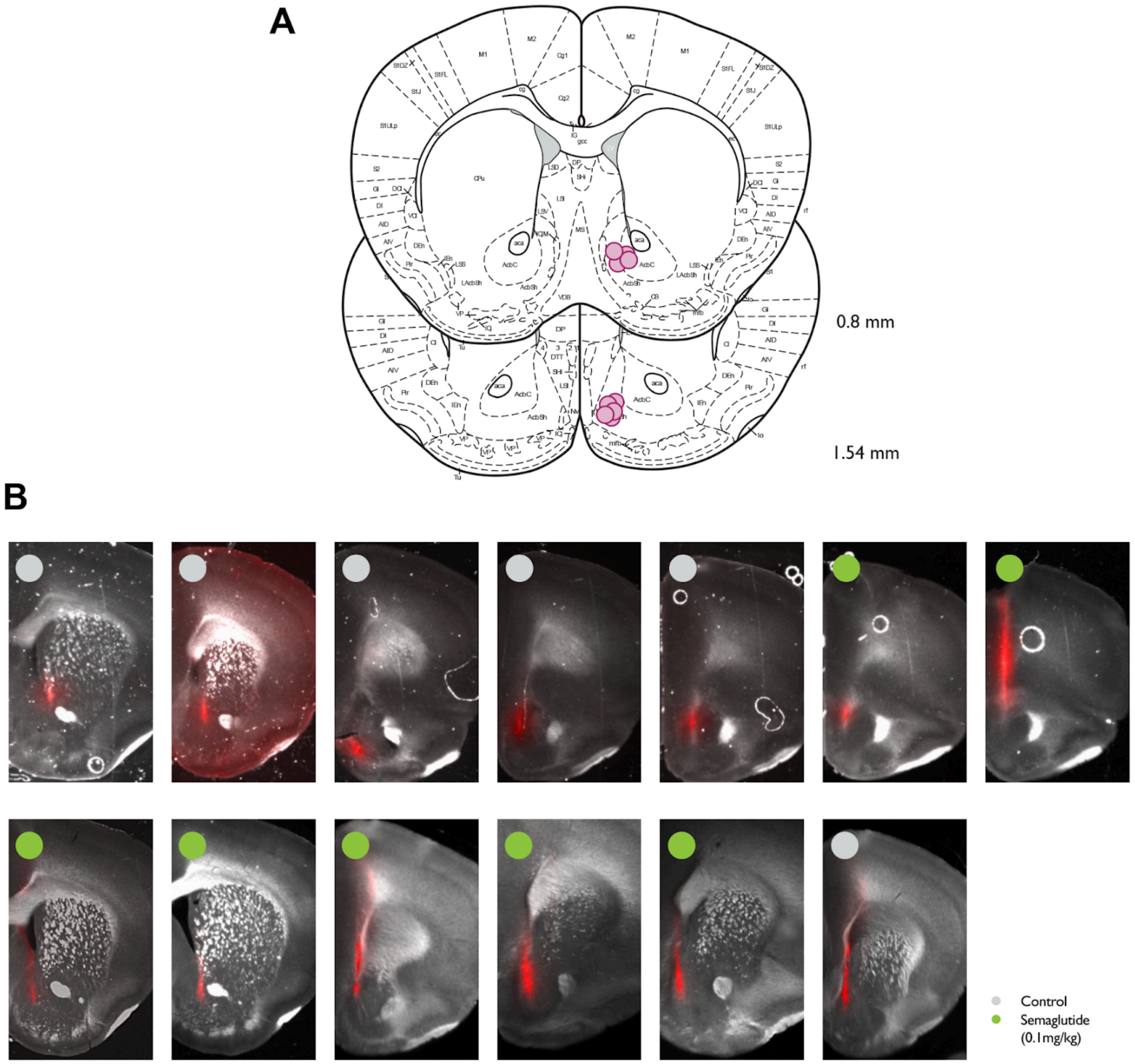


**Supplementary Figure 3. Histological analysis of recording electrode locations.** (A) Schematic from a standard brain atlas illustrating the estimated placement of the recording electrodes. (B) Representative histological sections from each animal, showing grayscale anatomical images overlaid with fluorescent DiI staining left by the electrode tip. This approach confirms that the electrode tracts align with the targeted brain region.
